# Discovering loop conformational flexibility in T4 lysozyme mutants through artificial intelligence aided molecular dynamics

**DOI:** 10.1101/2020.04.08.032748

**Authors:** Zachary Smith, Pavan Ravindra, Yihang Wang, Rory Cooley, Pratyush Tiwary

## Abstract

Proteins sample a variety of conformations distinct from their crystal structure. These structures, their propensities, and pathways for moving between them contain enormous information about protein function that is hidden from a purely structural perspective. Molecular dynamics simulations can uncover these higher energy states but often at a prohibitively high computational cost. Here we apply our recent statistical mechanics and artificial intelligence based molecular dynamics framework for enhanced sampling of protein loops in three mutants of the protein T4 lysozyme. We are able to correctly rank these according to the stability of their excited state. By analyzing reaction coordinates, we also obtain crucial insight into why these specific perturbations in sequence space lead to tremendous variations in conformational flexibility. Our framework thus allows accurate comparison of loop conformation populations with minimal prior human bias, and should be directly applicable to a range of macromolecules in biology, chemistry and beyond.

## Introduction

Understanding and predicting the relationship between protein sequence, structure and function has been a long-standing dream in biophysics. One of the many reasons why this is an especially difficult problem is that often a given sequence does not imply one fixed structure. On one hand, the so-called “folding problem” can be considered solved either through experiments (*1–4*) or even artificial intelligence (AI) (*5*). On the other hand, while it is useful to consider the single most stable protein crystal structure for a given sequence determined through experiments and/or theory, the typical model of proteins has shifted from static objects to fluctuating polymers (*6–8*). Considering the fluctuations of proteins between various structures, even rare ones, has been shown to increase our understanding of the mechanisms underlying protein function (*9–11*), yet these fluctuations are hard to quantify through even state-of-the art experiments (*11–13*). One particularly interesting class of conformational rearrangements is the movement of surface-exposed loops which exhibit much greater flexibility than the rest of the protein. This class of problems is relevant both from the perspective of a fundamental understanding of the chemistry of life processes (*14, 15*), as well as discovering druggable targets for inhibiting different diseases that offer both specific and potent action (*16, 17*).

A particularly well-characterized yet puzzling system for studying loop movement as well as the related problem of ligand recognition is the L99A mutant of the protein T4 lysozyme, in which a leucine residue at position 99 is replaced with alanine, opening up a pocket for binding small hydrophobic ligands such as benzene (*18*). This L99A mutated protein populates two well-defined states: the ground state (GS) and an excited state (ES), with populations of roughly 97% and 3% respectively at 25°C (*19*). The excited state is characterized by the rotation of the phenylalanine residue F114 into the binding pocket and unification of two adjacent helices (*20*). Additionally, benzene egress appears to take place during the transition from the ground state to the excited state, as the F114 moves into a buried position occupying the cavity previously filled by benzene (*21, 22*).

Previous computational studies have attempted to detail underlying mechanisms of this transition, but many aspects are still not fully understood (*23, 24*). For instance, in addition to the L99A mutant, two other point mutants of T4 lysozyme have been discovered which very significantly alter the population of the ground and excited states (*20*). These two mutants, L99A, G113A (double mutant) and L99A, G113A, R119P (triple mutant) have 66:34 and 4:96 ground state:excited state population ratios as opposed to the 97:3 for the single mutant L99A (*20*). A detailed atomic level insight into why and how such point mutations (Fig. 1) so drastically affect this protein’s conformational landscape is still missing. While computational and experimental methods have clarified such structural fluctuations and ligand binding pathways in L99A T4L in recent years, this system still presents a challenge due to the high degree of flexibility and size of the surface-exposed loop near the binding pocket (*23, 25*). This set of three mutants thus serves as an excellent benchmarking set for methods that seek to sample loop conformations including this current work.

**Figure 1:**
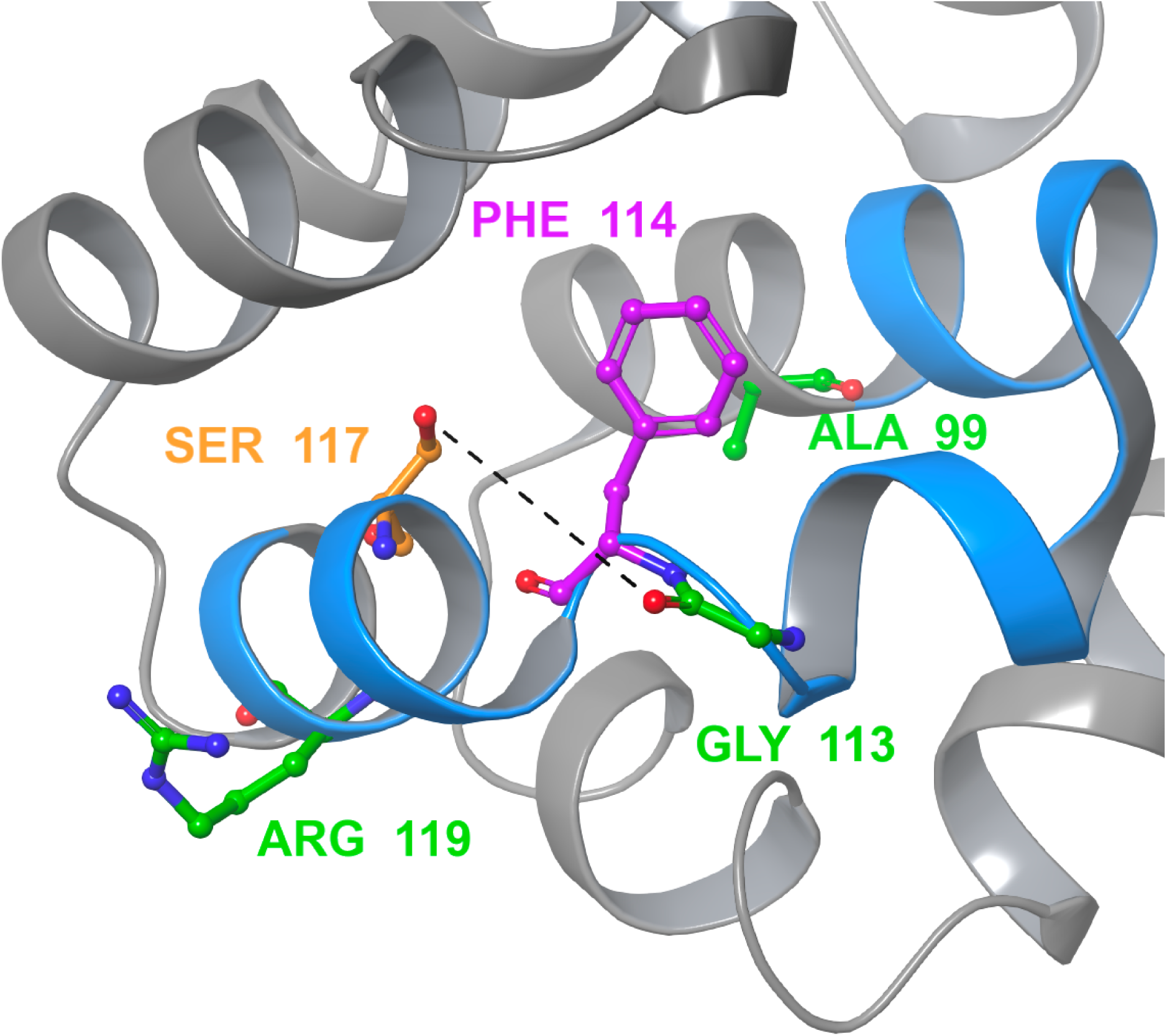
The crystal structure of the single mutant L99A with the cavity-adjacent loop (residues 100-120) shown in blue. The mutation sites for the three mutants are shown in green. The phenylalanine residue that blocks the binding pocket in the excited state is shown in purple. The distance between serine 117 (orange) and glycine 113 (green) that represents the formation of the hydrogen bond connecting the phenylalanine-adjacent helices is shown as a dashed line.

Our aim in this work is to demonstrate how our recent enhanced molecular dynamics (MD) simulation algorithms grounded in statistical mechanics and AI (*26–30*) can be used to obtain such insight in this classic test-piece system, thereby opening the possibility to answer such questions in the future in generic proteins with related unanswered questions of practical and fundamental relevance (*31, 32*). In principle, MD simulations can give such insights into the thermodynamics and biophysical mechanisms behind these protein conformational changes by providing data at the all-atom and femtosecond resolution, which can be expensive to obtain in experiments (*33*). However, MD simulations have a crippling timescale problem: reaching even the millisecond threshold on even the most specialized supercomputers is very difficult due to the sheer number of interactions that need to be computed per timestep and the short timescale of bond vibrations restricting each timestep to a femtosecond or two at best (*34*). Often, these metastable protein conformations are separated by high energy barriers that make exchanges between these conformations much slower than hundreds of microseconds and therefore practically inaccessible to classical MD simulations (*28*).

To overcome this timescale problem, enhanced sampling methods have been designed to accelerate the sampling of protein configurations (*35–37*). One such method is metadynamics (*28, 38, 39*). If given a carefully constructed reaction coordinate (RC) that has sufficient overlap with the relevant slow degrees of freedom, metadynamics can gradually enhance fluctuations along this RC to efficiently move the system between known and unknown metastable states in a controlled, reweightable manner (*28*). The major limitation of metadynamics and arguably many other enhanced sampling methods as well is the dependence of its performance on the selection of the RC whose fluctuations are enhanced through biasing. Traditionally, this RC has been chosen by hand using chemical intuition and results from previous studies. But in recent years methods have been developed which aim to learn RCs with minimal human intuition (*26, 27, 40–48*).

One such AI-inspired method is Reweighted Autoencoded Variational bayes for Enhanced sampling (RAVE), which is based on the principle of predictive information bottleneck (PIB), originally proposed to model neuronal behavior in retinal cells (*49*) and more recently as an explanation for the generic success of deep learning and AI (*50*). In a nutshell, the PIB is the minimally complex yet maximally predictive model for describing the evolution of a given dynamical system (*27, 51*). RAVE learns such a PIB in an iterative procedure, interpreting it as the RC to be used as a biasing variable in enhanced sampling (*26, 27*). Here, we construct this RC as a linear combination of selected basis functions or order parameters (OPs). These OPs can number in the hundreds to thousands and are generic features such as protein-ligand or protein-protein contacts. Here we first use our method Automatic Mutual Information Noise Omission (AMINO) (*30*) that screens for redundancies among this very large and generic set of OPs, screening for example if two protein-ligand contacts carry the same information or not. The output from AMINO is fed to RAVE (*26, 27*) which then uses the PIB to construct a RC as a linear combination of the AMINO OPs. The RC is then used as biasing variable in metadynamics, and the biased metadynamics trajectory itself is fed back to RAVE to further optimize the RC. The iteration between deep learning and sampling continues until multiple transitions between different metastable states are sampled. While this study uses metadynamics, any enhanced sampling method that uses a biasing RC can be used with AMINO and RAVE. Through this automated protocol, we are able to compare the changes in ground and excited state populations due to point mutations and gain insights into the underlying mechanisms causing these changes. We are also able to analyze the RC and its constituent basis functions for different sequences and gain mechanistic insight into how small perturbations in sequence space lead to enormous effects in the structure space. We expect that the procedure using AMINO, RAVE, and enhanced sampling illustrated in this work can be generalized to sample a wide array processes undergone by macromolecules, especially those which have not been studied extensively enough to have standard enhanced sampling parameters.

## Results

### Summary of Methods

Here we briefly summarize the methods used in this work, with full details given in **Materials and Methods**. Short unbiased MD simulations were performed for each of the three mutants starting from their respective x-ray crystal structures. Starting from a much larger list of different inter-residue contacts, AMINO (*30*) generated a minimal, representative set of OPs for each of the 3 systems. A first round of RAVE was then applied to the unbiased MD trajectory to learn a RC as a linear combination of the OPs output from AMINO. Well-tempered metadynamics (*28*) simulations were performed using this RC to accelerate the sampling, and the resultant biased trajectory was fed back to RAVE to learn an improved RC. This iterative procedure allows us to explore conformational space much faster than traditional MD while returning RCs in terms of human-interpretable variables such as the distances between C*α* atoms used in this study. The iteration between RC optimization through RAVE and sampling through well-tempered metadynamics was terminated when the latter achieved multiple back-and-forth transitions between the starting state and other metastable states. Once RC optimization was completed, two longer 500 ns productions runs of metadynamics were performed for each mutant.

### Order Parameter and Reaction Coordinate Analysis

The minimal OP set output from AMINO and the subsequently optimized RCs are shown for the 3 different mutants in Table 1. Two comments are critical here. First, as can be seen in Table 1, the dimensionality of the minimal OP set differed for the three mutants. Second, the RCs were all 2-dimensional. We first attempted to perform the protocol of this work with a 1-dimensional RC, but subsequent rounds of metadynamics and RAVE were found to show no improvement in sampling. As such we used a 2-dimensional RC wherein sampling improved with subsequent training rounds. This RC was comprised of components *χ*_1_ and *χ*_2_ which themselves were expressed as linear combinations of the input OPs.

**Table 1:** Reaction coordinate definitions for each mutant. *χ*_**1**_ and *χ*_**2**_ refer to the first and second component of a two-dimensional RC. The order parameters are defined as the distance between two C*α* atoms. Order parameters using two cavity-adjacent loop residues are shown before other order parameters and are in italics.

| **Single Mutant Distances** | $\chi_1$ Weight | $\chi_2$ Weight | **Double Mutant Distances** | $\chi_1$ Weight | $\chi_2$ Weight | **Triple Mutant Distances** | $\chi_1$ Weight | $\chi_2$ Weight |
| --- | --- | --- | --- | --- | --- | --- | --- | --- |
| *G110 - L118* | -0.249 | 0.895 | *G110 - V111* | 0.820 | 1.000 | *V111 - A112* | -0.047 | -0.083 |
| *V111 - A112* | -0.498 | 0.052 | *A112 - A113* | 0.660 | 0.110 | *N116 - S117* | -0.248 | -0.103 |
| *A112 - G113* | 0.221 | -0.017 | *A113 - S117* | 0.307 | 0.326 | *S117 - L118* | 0.087 | 0.559 |
| *T115 - N116* | -0.197 | -0.075 | *N116 - M120* | -0.262 | -0.228 | I27 - T115 | 1.000 | -0.547 |
| Y25 - T115 | -0.056 | -0.406 | *S117 - L118* | -0.715 | -0.430 | F114 - T152 | 0.476 | 1.000 |
| G28 - G110 | 0.083 | -0.411 | *L118 - R119* | -0.651 | -0.298 |  |  |  |
| T34 - F114 | -0.409 | -0.006 | K43 - V111 | 0.256 | -0.099 |  |  |  |
| T34 - L118 | 0.669 | 0.385 | R76 - V111 | 0.558 | -0.181 |  |  |  |
| K43 - F114 | 0.120 | -0.425 | R119 - K124 | -0.133 | 0.435 |  |  |  |
| A49 - N116 | 0.062 | -0.162 | M120 - L121 | 1.000 | 0.256 |  |  |  |
| F67 - N116 | 0.237 | 0.767 |  |  |  |  |  |  |
| K85 - A112 | -0.016 | 0.297 |  |  |  |  |  |  |
| A97 - T115 | 1.000 | -0.701 |  |  |  |  |  |  |
| T142 - R119 | 0.206 | 1.000 |  |  |  |  |  |  |
| R154 - G113 | 0.082 | 0.393 |  |  |  |  |  |  |
| T157 - N116 | -0.056 | 0.975 |  |  |  |  |  |  |

Looking at the dimensionality of the minimal OP set for each mutant we can determine how many independent components are needed to describe the loop’s dynamics and what type of components are needed to do so. We can see that the single, double and triple mutants have 16, 10 and 5 independent OPs respectively. Thus for the single mutant, there are more possible contacts whose fluctuations can lead to change in the RC, or equivalently, whose fluctuations can cause back and forth movement between metastable states. Moreover, each subsequent mutation appears to lower the number of OPs or equivalently lowers the global flexibility of the protein. While the single mutant needs an overall higher number of contacts to describe the RC for conformational change, even more interestingly, the doube mutant has a larger number of these contacts between residues from the loop region. This means that the double mutant’s loop conformational rearrangements have a greater number of degrees of freedom, or in other words, the loop in the double mutant is more flexible than the other two mutants while the single mutant exhibits greater global flexibility (*52*).

In general the use of AI-based frameworks suffer from an interpretability issue due to the black-box nature of AI. In RAVE the use of a simple linear encoder circumvents this problem and allows us to directly look at the weights of the OPs themselves. When considering the highest three weights for each RC component of each mutant (shown in Fig. 2), the single mutant has few inter-loop contacts, while the double mutant has primarily inter-loop contacts and the triple mutant has a mix of inter-loop contacts. This again means that not only does the double mutant show greater local flexibility but that this local flexibility plays a key role in the cavity-adjacent loop’s rearrangements, while the triple mutant is globally as well as locally less flexible than the other two given the much smaller number of independent OPs. Further, we needed to consider a two-component RC to sample the loop’s movement which implies that there are two processes that must occur in order for the conformation to change.

**Figure 2:**
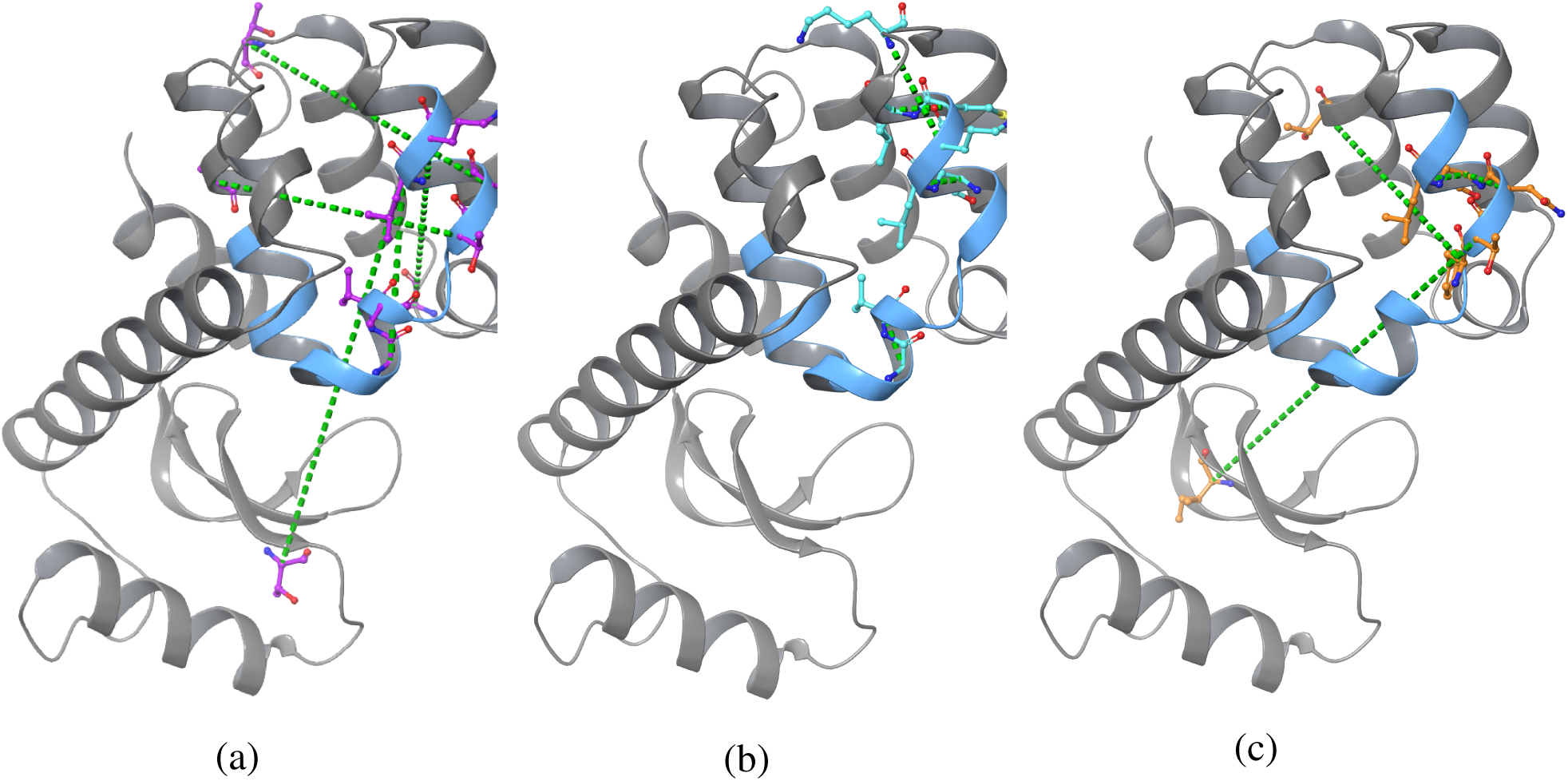
Visual representation of the three highest weighted OPs in each of the two RC components for each mutant. The OPs are shown as dashed green lines while the cavity-adjacent loop is highlighted in blue. The residues used in OPs are shown in purple for the single mutant (a), cyan for the double mutant (b), and orange for the triple mutant (c). There are fewer than 6 OPs shown for (b) and (c) because some OPs were among the highest three weights for both RC components.

In summary so far, we see from analyzing the RCs as well as the OPs needed to build them, that subsequent mutations lower the protein’s global flexibility, and from the increased number of inter-loop OPs that the double mutant’s loop residues have increased local flexibility. While conformational change in the single and the triple mutants needs to be triggered by long-distance fluctuations in the protein, changes in the double mutant can be triggered through short-distance fluctuations. This difference could possibly be a reason for why these three sequences differ so distinctly in their propensity for taking different conformations.

### Free Energy Calculation

With the two-dimensional RCs as described in the previous sub-section for different mutants, we then performed 500 ns of metadynamics for each mutant. Two independent copies were performed to reduce noise and ascertain run-to-run variability. The effect of the metadynamics bias was reweighted in order to recover unbiased statistics (*53*). These reweighted statistics were then combined with definitions of various states/conformations to calculate the free energy difference between the ground and excited states. To stay consistent with literature (*23, 24*) we defined these states in terms of three variables: (i) the F114 Ψ dihedral angle, (ii) the hydrogen bonding distance between G113 and S117, and (iii) the RMSD deviation over C*α* atoms from the non-native conformation for each mutant (PDB: 2LCB for single and double mutants and PDB: 1L90 for the triple mutant). The distance and dihedral angle shown in Fig. 1 have been used in previous studies (*23*) but we found them to be insufficient to define the states as they could yield false positives. Specifically, the secondary structure of the cavity-adjacent loop may become distorted over time which may yield conformations that match the distance and dihedral criteria for a state but do not correspond to the entry and exit of the cavity by F114. Thus in order to avoid getting trapped in these conformations we only considered conformations with RMSD below a given threshold.

These unfolded states must be accounted for in both state definition and metadynamics simulation. Even a biased simulation with an optimized RC, or equivalently a very long unbiased simulation, can get trapped in these unfolded configurations with very high RMSD from any state of interest. In principle these trapping states are not a problem as the simulation would eventually return to regions of greater interest. However, they can lead to debilitating computational efficiency. Therefore, we included a wall in terms of RMSD that prevents the protein from exploring conformations far from the states of interest in our metadynamics simulation (see details in **Materials and Methods**). The state definitions using distance, dihedral, and RMSD were set using a semi-automatic procedure whose details are provided in **Materials and Methods**.

Once these state definitions, shown in Table. 2, had been established for each mutant, we calculated the relative thermodynamic stability of these conformations, as quantified by their free energy difference. In Fig. 3 we show the free energy differences between the excited and ground states over the two independent 500 ns metadynamics runs each for the single, double and triple mutants respectively. While there is run to run scatter, the averaged values shown in Fig. 4 (see **Materials and Methods** for averaging and error estimate protocols) are in excellent qualitative and quantitative agreement with the experimental values reported by Bouvignies et al. (*20*). It should be noted that the plateaus in free energy in Fig. 3 can correspond to either a converged estimate of the free energy difference or prolonged exploration of conformations that do not correspond to their the ground or excited state. However, as can be seen from the trajectories of different OPs as well as movies provided in Supplemental Information (SI) we have ample movement between different conformations. In the SI we also provide various free energy profiles (Figs. S1-S4) along relevant OPs, including the RMSD, and RC components. These profiles show clearly as to why we needed to use restraints in the RMSD space. In order to ascertain how sensitive our calculations are to the use of such restraints, we did not apply such a restraint for the triple mutant. While we still obtained relative free energy in good agreement with experiment (Fig. 3), the corresponding Fig. S3 in SI when compared with Figs. S1 and S2 where we did use restraints shows the usefulness of doing so. Finally, Fig. S4 in the SI shows how the estimate from Fig. 3 would have changed with different RMSD values used in defining states (Table 2). The values are extremely robust until we reach the range where the restraints were active and the sampling is no longer reliable.

**Table 2:** State definitions for the ground and excited states of each mutant using F114 Ψ, 113-117 distance, and RMSD.

| **Mutant** | **GS Distance (Å)** | **GS $\Psi$ (radians)** | **ES Distance (Å)** | **ES $\Psi$ (radians)** | **RMSD Cutoff (Å)** |
| --- | --- | --- | --- | --- | --- |
| L99A | $7 \pm 1.5$ | $.75 \pm .375$ | $2.5 \pm 1.5$ | $-.75 \pm .375$ | $< 3.6$ |
| L99A, G113A | $6 \pm 1.5$ | $.75 \pm .375$ | $2.5 \pm 1.5$ | $-.30 \pm .375$ | $< 4.4$ |
| L99A, G113A, R119P | $6 \pm 1.5$ | $.75 \pm .375$ | $2.5 \pm 1.5$ | $-.75 \pm .375$ | $< 5.0$ |

**Figure 3:**
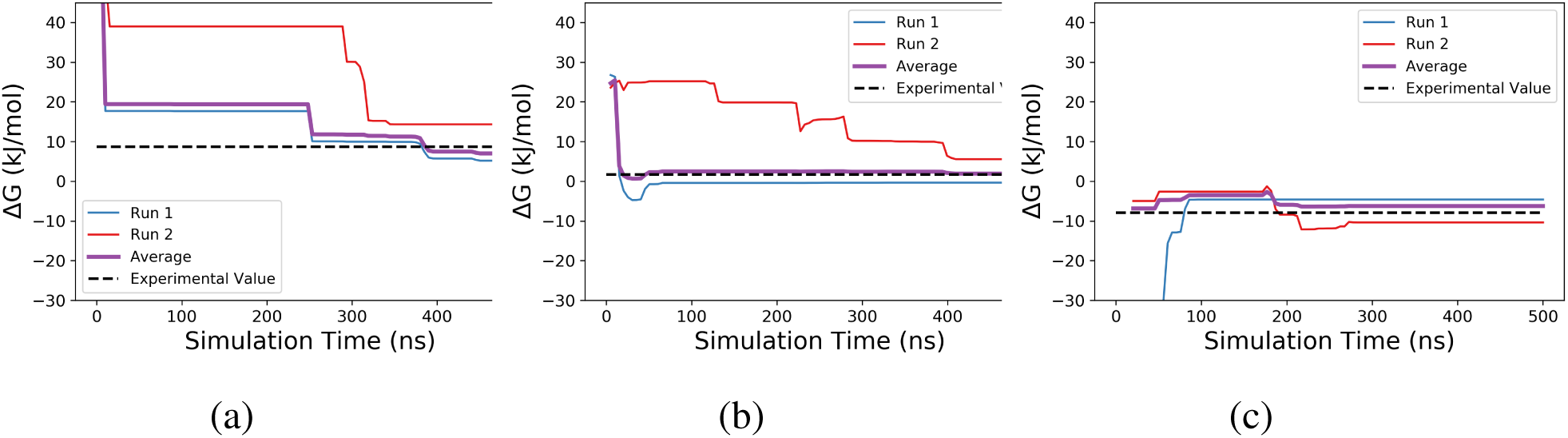
Free energy differences between the excited and ground states over time for the single mutant (a), double mutant (b), and triple mutant (c) starting from the first timestep where both the ground and excited states have been visited. The evolution of ΔG for each replica is shown in red and blue. The ΔG corresponding to the average probabilities of the two replicas is shown in purple. The ΔG value reported in 20 is shown as a dashed line.

**Figure 4:**
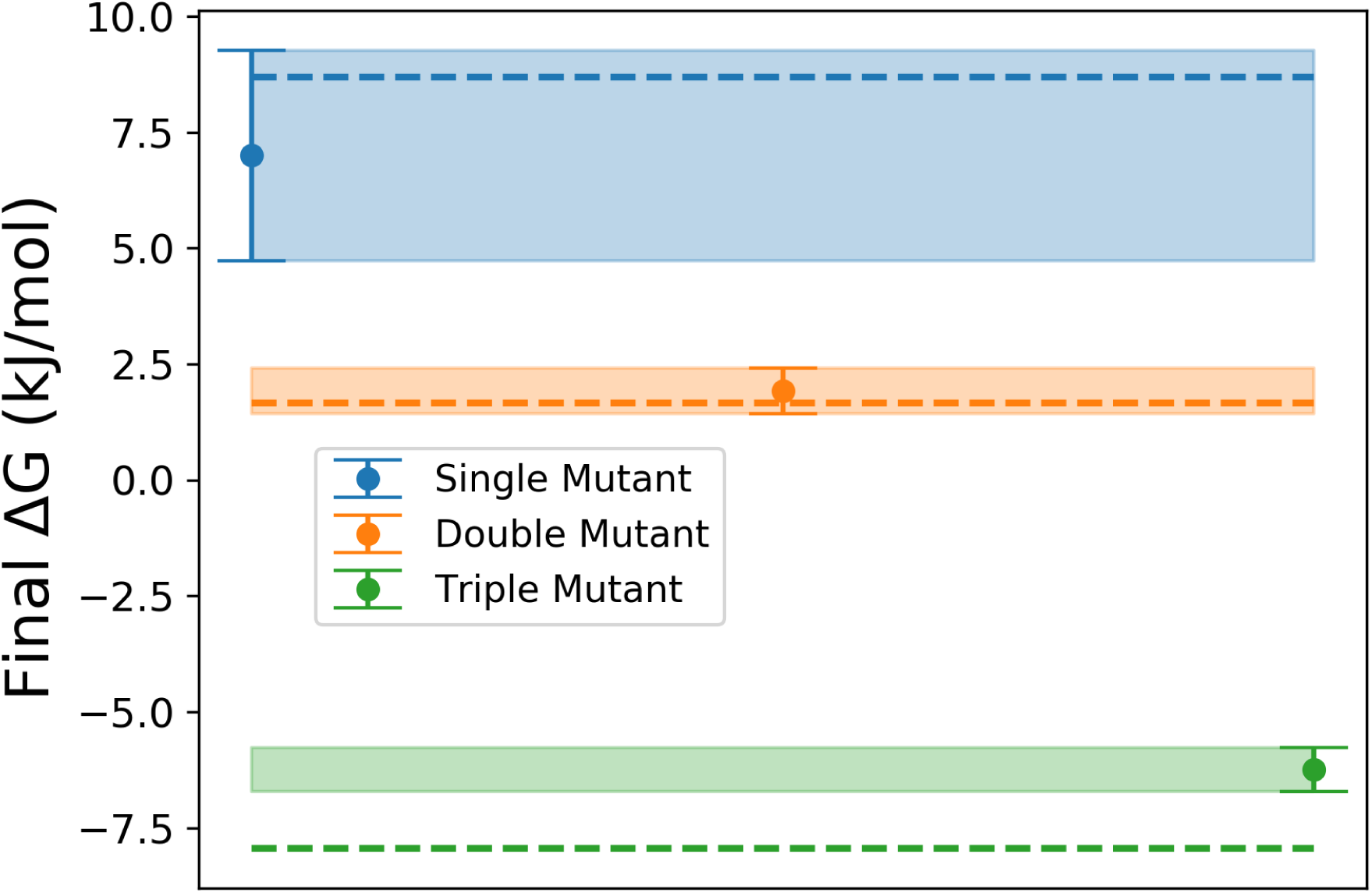
Free energy differences between the excited and ground states after 500 ns of metadynamics simulation was performed using two replicas for each state. Each mutant is represented by the final ΔG value with error bars obtained by block averaging. For comparison, the ΔG corresponding to the distributions reported in Ref. 20 are shown as dotted lines matching the color of the corresponding mutant.

## Discussion

In this work we have proposed and demonstrated an all-atom molecular dynamics based simulation protocol for quantifying flexible loop conformational propensities as a function of the underlying protein sequence. The enhanced sampling procedure of this work represents an automated framework for sampling loop conformations which can be used for both free energy ranking of known states and discovery of new states. Here we have applied to the classic but challenging test-piece of mutations in the T4 lysozyme family of proteins. Comparing the final free energy differences and their error calculated with block averaging (Fig. 4), a clear ranking emerges recovering the correct order of thermodynamic stability. Not only is the ranking correct, but for all 3 mutants we obtain conformational free energy differences that are nearly identical to the experimental values, within thermal fluctuation and sampling error margins. In addition to obtaining correct thermodynamic propensities of the different conformations, we have also gained crucial atomic-scale insight into the differences in the dynamics between the three mutants. We see that the total number of OPs needed to describe the dynamics of each mutant decreases as additional mutations occur suggesting a decrease in global flexibility. Despite the trend with the total number of OPs, the double mutant has the largest number of inter-loop OPs suggesting increased local flexibility in the cavity-adjacent loop. We also obtain insight from looking at the weights of the different OPs that build up the RC for the 3 systems. We see that the increased local flexibility in the double mutant plays a key role in its conformational rearrangements.

While this work presents a step towards a fully automated procedure for sampling loop conformations in all-atom resolution, we would like to highlight that the state definitions for loop conformations is still an open problem that requires a large quantity of expert knowledge. Though state definition is not yet automated in our work, thereby leading to us not calling it fully automated as of yet, progress has been made in developing generalized methods for state definition for example using path collective variables (*24*). In conclusion, the combination of statistical mechanics and AI-based methods AMINO, RAVE and metadynamics allowed us to enhanced the sampling of the ground and excited states in flexible proteins in a semi-automatic manner with minimal human intervention. Through this we could obtain accurate free energies as well as physically relevant basis functions that give direct mechanistic understanding into protein conformational plasticity, and hopefully inspire future experimental or computational studies.

### Supplementary Material

accompanies this paper at http://www.scienceadvances.org/.

## Materials and Methods

### AMINO

In this work, we constructed our RC as a linear combination of a set of basis functions, which we call order parameters (OPs). AMINO starts with a large dictionary of OPs and uses an information theoretic approach to identify redundancies in the full dictionary to return a reduced set of OPs for use in RC construction (*30*).

AMINO runs a k-medoids clustering procedure on the set of OPs based on the following distance metric:

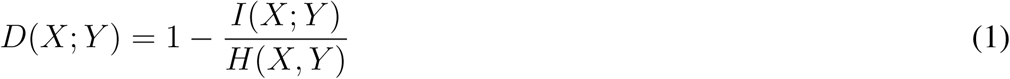

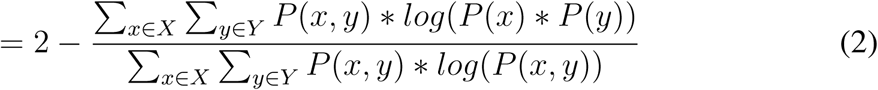

As input to AMINO for each system, we used a combination of a 50 nanosecond unbiased trajectory beginning in ES and a 50 nanosecond unbiased trajectory beginning in GS to form a combined trajectory of 100 nanoseconds. Since AMINO needs only estimates of the stationary probability density, preserving the temporal ordering of data points is not needed, and this kind of mixing is acceptable. From these trajectories the contact points used to construct the OPs for each system were the following:

1. Category I: The C*α* of every amino acid in the loop, which were defined to be amino acids in the range [110, 120].
2. Category II: Every third C*α* that is not in the above loop range.

In the second category in principle we could take every amino acid and not just the third, but the latter helps with computational efficiency. In future versions of AMINO we hope to make it possible to consider every amino acid in a computationally efficient manner. The input OPs to AMINO were the distances between every pair of contact points *p, q* where *p* is a contact point from Category I, *q* is a contact point from either Category I or II, and *p* ≠ *q*.

From this input data, AMINO runs the aforementioned clustering procedure on these OPs for a range of *k* values, where *k* is the number of clusters used in k-medoids clustering. AMINO decides the optimal value of *k* automatically as discussed in Ref. 30. This forms the procedure that allows for automated selection of OPs with minimal prior knowledge of the system.

### RAVE

RAVE took the output OPs from AMINO and uses a information bottleneck based protocol to learn the RC, which was then used in metadynamics to enhance the sampling. The sampling from metadynamics is then fed back to RAVE to learn a better RC, which in turn is used to perform a newer round of metadynamics. The iteration between metadynamics and RAVE is continued until sufficient back-and-forth movement between metastable states is obtained during metadynamics. In RAVE, the RC is defined as the predictive information bottleneck which encodes the high dimensional input *X* as a low dimensional representation *χ*, which we interpret as the RC. The optimally encoded RC should be the low dimensional representation that maximally compresses the input, yet also has maximal predictive power for the future state of the input, given by *X*_Δ*t*_. This trade-off between maximal compression and maximal prediction can be quantified using the two mutual informations *I*(*χ, X*) and *I*(*χ, X*_Δ*t*_), or more specifically, the difference between these two mutual informations (*27,29*). This has been shown to be equivalent to training a neural network with an encoder-decoder structure to optimize the following objective function (*54*):

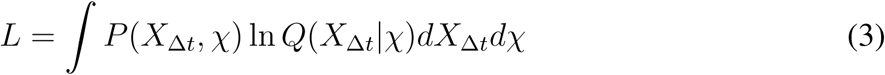

Here *χ* is a linear combination of OPs determined by the encoder and *Q*(*X*_Δ*t*_*|χ*) is the probability learned by the deep neural network to approximate the distribution *P* (*X*_Δ*t*_*|χ*) from data. In this study, we needed to iterate between metadynamics and RAVE so we needed to reweight the effect of biasing potential on the static probability as well as the dynamical propagator. The correction introduced in Ref. (*29*) is used to give a better estimation of *P* (*X*_Δ*t*_, *χ*) from biased trajectories.

### Molecular Dynamics

All molecular dynamics simulations were run using the AMBER99SB force field through the GROMACS package patched with PLUMED ver 2.4.2 (*55,56*) with a 2 fs timestep. The starting structures were PDB: 1L90 for the single mutant, PDB: 3DMV for the double mutant, and PDB: 2LC9 for the triple mutant. The triple mutant was prepared by mutating residue G113 to A113 using MAESTRO on the starting structure.

The AMINO and RAVE inputs were 50 ns of simulation from the starting structure and 50 ns of simulation from the less-stable state (ES for single and double mutants and GS for triple). The starting structures for the less stable states were set by finding the minimum RMSD between a reference structure for that state and a metadynamics trajectory along a trial inter-residue contact RC that visited both states. This RMSD was calculated by aligning all C*α* except 100-120 and measuring the displacement of C*α* 100-120. The excited state reference structures were PDB: 2LCB for single and double mutants and PDB: 1L90 for the triple mutant. Mixing these two trajectories for each system allowed us to consider the fluctuations in both states when constructing a basis set of OPs and a RC.

### Metadynamics

The metadynamics simulations were run using the PLUMED implementation of well-tempered metadynamics with biasfactor of 10, intitial hill height of 1.5 kJ/mol, and bias deposited every ps (*39, 55, 56*). The sigma value of the gaussian bias kernel was set to the standard deviation of the biasing RC from 50 ns of unbiased simulation for each system.

Once an initial 15 ns metadynamics run was completed for each system, the biased trajectory was reweighted and used to run a second round of RAVE (*57, 58*). The system in these 15 ns explored more conformational space than the initial 100 ns of unbiased simulation which enabled RAVE to determine a more informative RC. The RC from the second round of RAVE was used for the longer production runs of metadynamics.

The production runs also included a quadratic bias for the single and double mutants to prevent the loop’s helix from breaking as bias accumulated. The quadratic bias (kJ/mol) had the form of Eq. 4 for the single mutant and Eq. 5 for the double mutant where RMSD (nm) is calculated using the protocol from the AMINO inputs. These metadynamics runs were performed in duplicate with randomized starting velocities for 500 ns for each of the three mutants.

The results for the triple mutant could be improved by adding a quadratic wall similar to those used for the other two mutants. The triple mutant spent a large quantity of time in high RMSD conformations that were not considered in the GS or ES which can be alleviated by adding such a barrier. We have chosen to leave one mutant without a barrier in this work to clearly show the difference between trajectories with and without a barrier. The use of these RMSD barriers makes the procedure more efficient but less automated than using AMINO and RAVE alone. This can be improved by the use of a metric that does not rely on crystal structures but describes secondary structure such as the alpha helix and beta sheet RMSDs developed by Pietrucci and Laio (*59*).

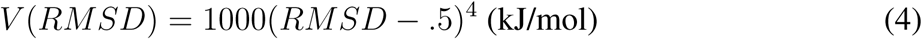

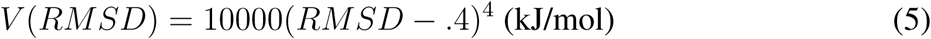

### State Definitions

The distance and Ψ dihedral ranges for each state were set using the equilibrium values along each variable for each mutant, where the equilibrium values were obtained from the average probabilities across the mutant’s two 500 ns metadynamics runs. The center of the ranges were set to the local minima closest to those reported in Ref. 23 and the range around these centers was set by hand to be a static range across mutants which covered the minima but overlap with the other state for any mutants. The RMSD cutoffs were then set to be the RMSD of the each mutant’s equilibrated structure + .1 Å.

## Acknowledgements

We thank Schrodinger LLC for providing us access to their software suite.

## Author Contributions

All authors contributed to running the simulations, analyzing the results, and writing the manuscript.

## Data and materials availability

The input files necessary for reproducing the metadynamics simulations reported in this paper can be found on PLUMED-NEST at plumed-nest.org/eggs/20/007/ (*56*). The scripts necessary for reproducing the OP and RC calculations can be found on GitHub at github.com/tiwarylab.

## Supplementary Material

**Movies**

1. Single mutant trajectory over 500 ns of metadynamics simulation. The cavity-adjacent loop is highlighted as blue while residue 114 is highlighted in magenta, available via Google Drive at https://tinyurl.com/T4-single
2. Double mutant trajectory over 500 ns of metadynamics simulation. The cavity-adjacent loop is highlighted as blue while residue 114 is highlighted in magenta, available via Google Drive at https://tinyurl.com/T4-double
3. Triple mutant trajectory over 500 ns of metadynamics simulation. The cavity-adjacent loop is highlighted as blue while residue 114 is highlighted in magenta, available via Google Drive at https://tinyurl.com/T4-triple

**FIG. 1:**
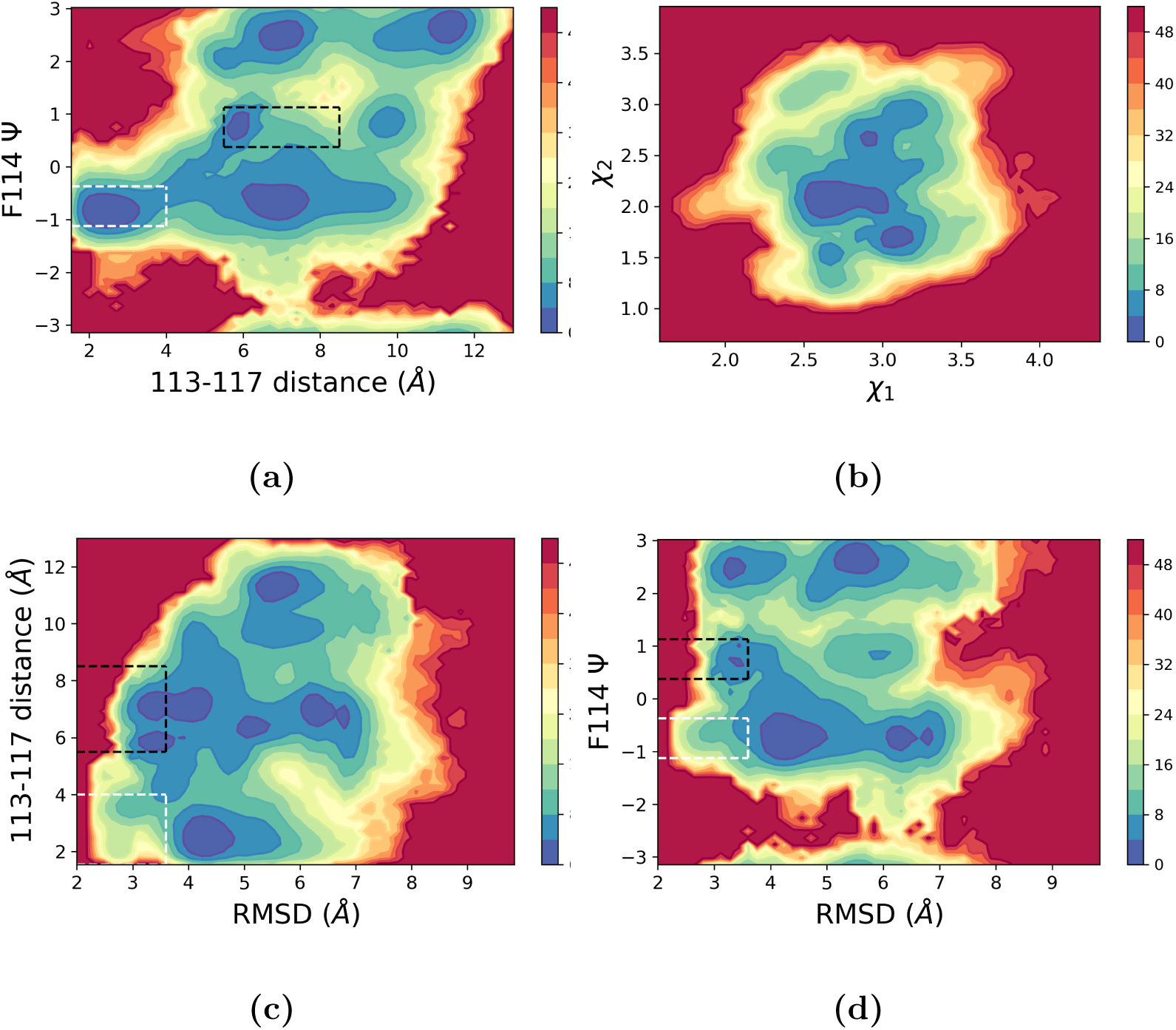
The single mutant free energy surface in kJ/mol projected along (a) the distance between c*α* 113 and c*α* 117, and the F114 Ψ dihedral angle, (b) the two reaction coordinate components, (c) RMSD and the 113-117 distance, and (d) RMSD and F114 Ψ. The ground state is shown in a dashed black box while the excited state is shown in a dashed white box. (b) displays the need for a 2D reaction coordinate as there are free energy wells that need both *χ*_1_ and *χ*_2_ to be distinguished from each other. (c) and (d) demonstrate the necessity for a RMSD cutoff by showing the change in state populations as RMSD increases to levels far from the native conformations.

**FIG. 2:**
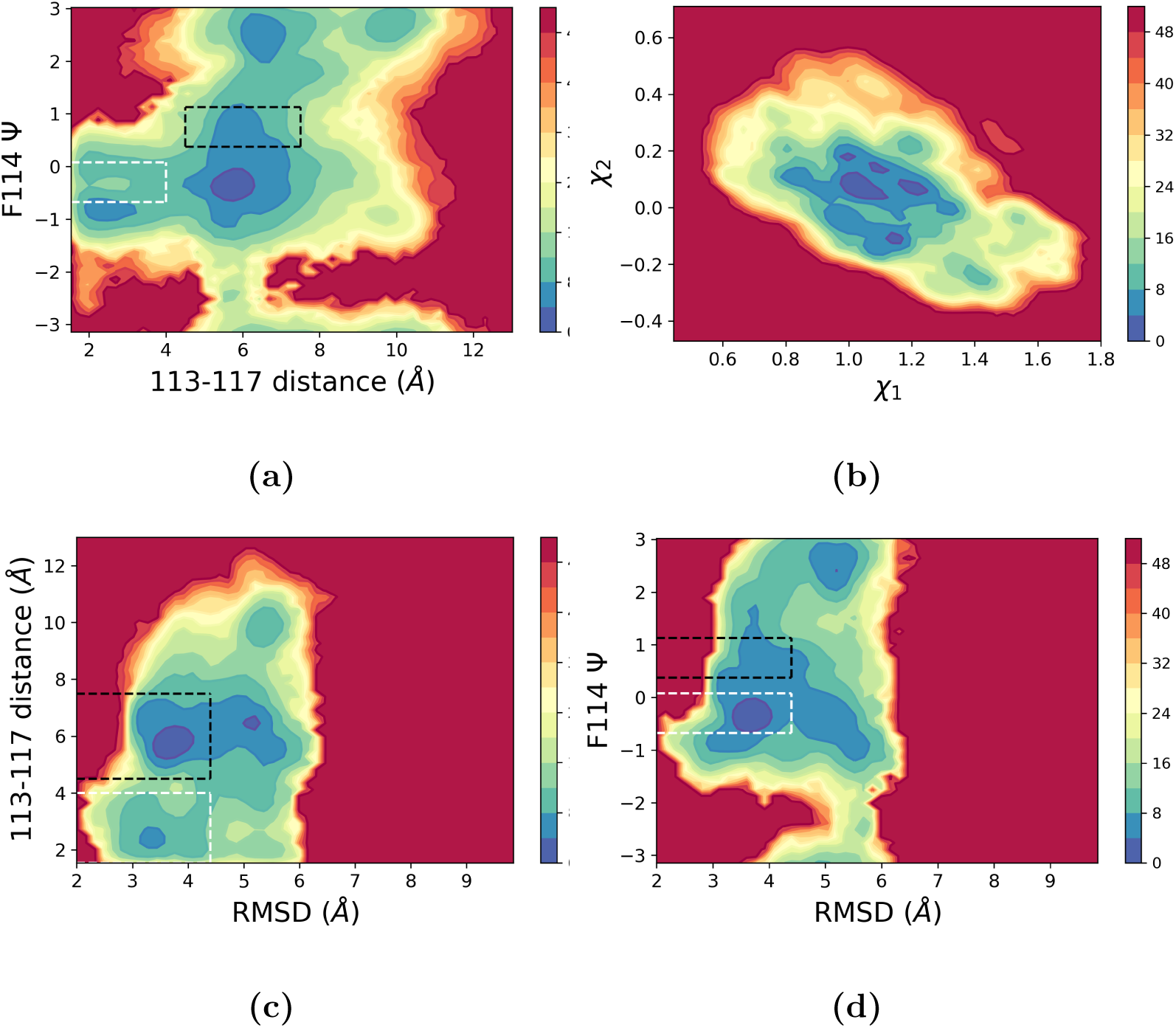
The double mutant free energy surface in kJ/mol projected along (a) the distance between c*α* 113 and c*α* 117, and the F114 Ψ dihedral angle, (b) the two reaction coordinate components, (c) RMSD and the 113-117 distance, and (d) RMSD and F114 Ψ. The ground state is shown in a dashed black box while the excited state is shown in a dashed white box. (b) displays the need for a 2D reaction coordinate as there are free energy wells that need both *χ*_1_ and *χ*_2_ to be distinguished from each other. (c) and (d) demonstrate the necessity for a RMSD cutoff by showing the change in state populations as RMSD increases to levels far from the native conformations.

**FIG. 3:**
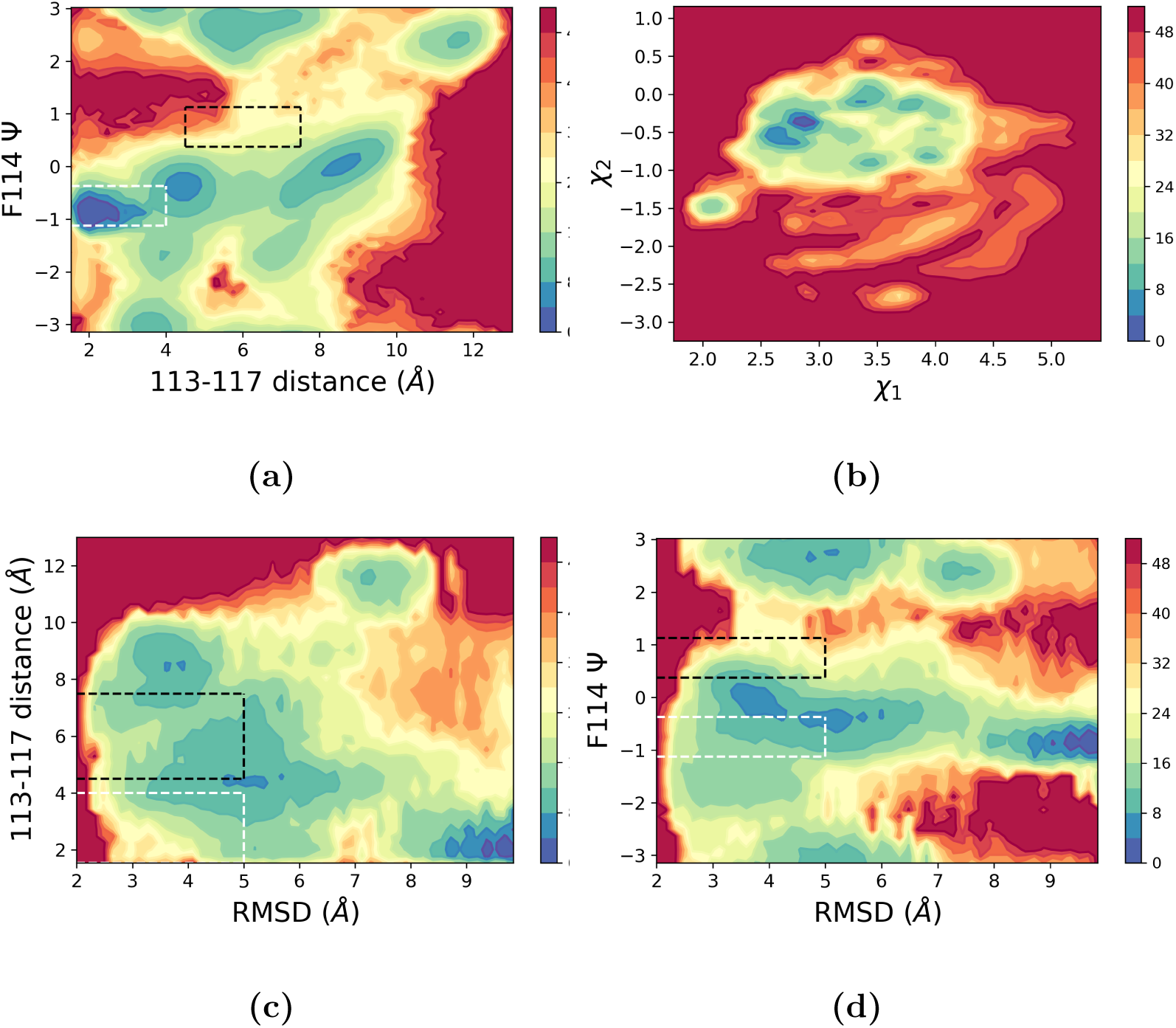
The triple mutant free energy surface in kJ/mol projected along (a) the distance between c*α* 113 and c*α* 117, and the F114 Ψ dihedral angle, (b) the two reaction coordinate components, (c) RMSD and the 113-117 distance, and (d) RMSD and F114 Ψ. The ground state is shown in a dashed black box while the excited state is shown in a dashed white box. (b) displays the need for a 2D reaction coordinate as there are free energy wells that need both *χ*_1_ and *χ*_2_ to be distinguished from each other. (c) and (d) demonstrate the necessity for a RMSD cutoff by showing the change in state populations as RMSD increases to levels far from the native conformations. Note that for this system we did not apply any RMSD restraints to see how robust our calculations are. This has led to an overall noisier free energy profile as the systems spends time in many trapping states described in the main text. However, as can be seen in the main text, we still obtain relative thermodynamic propensities for the two conformations in decent agreement with experimental benchmarks.

**FIG. 4:**
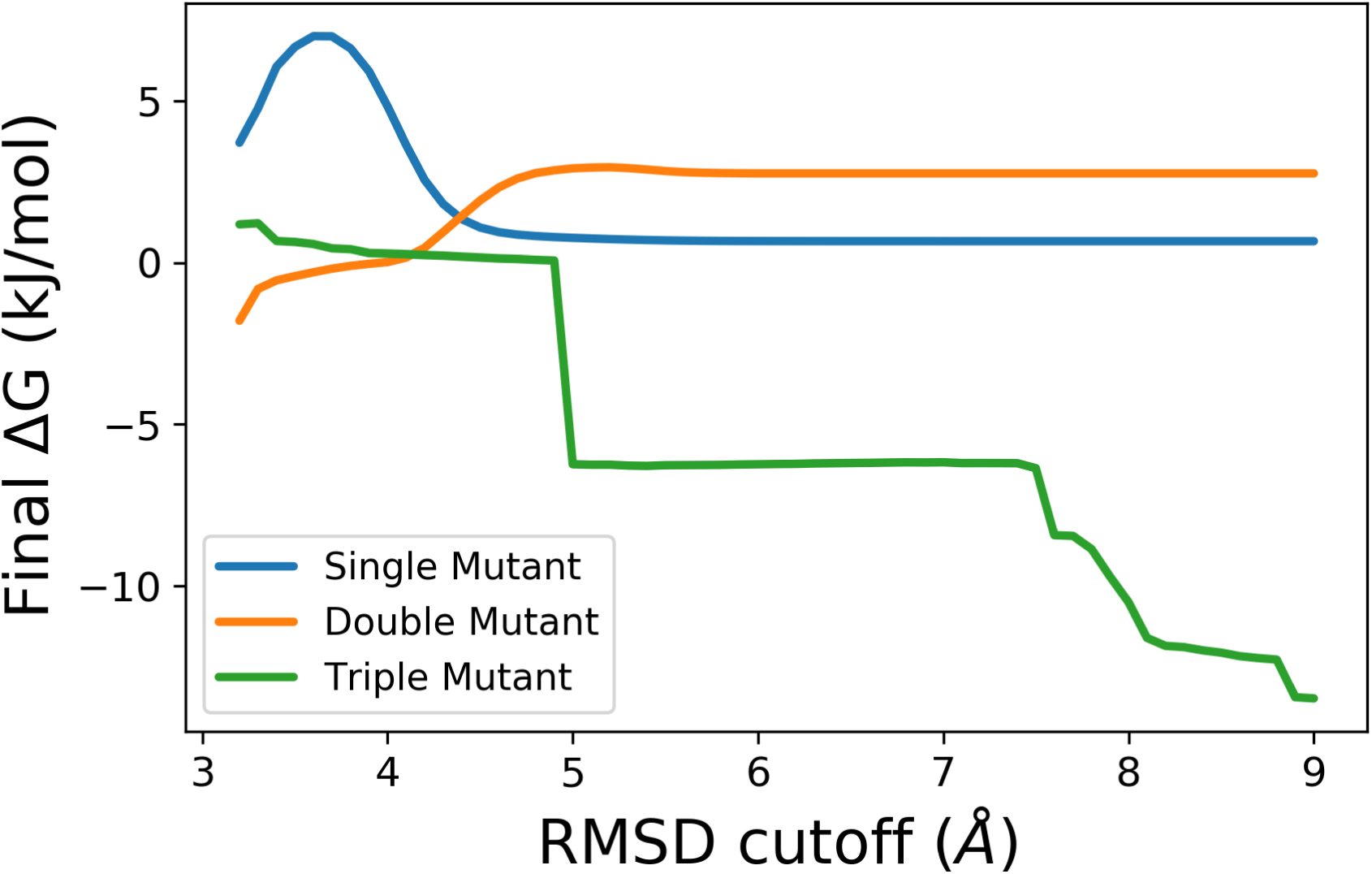
Final free energy differences for all three mutants calculated using varying RMSD cutoff values in the state definitions. With a misleading RMSD cutoff, we either exclude one of the states or include conformations where the cavity-adjacent loop is distorted. These effects can be seen in the spikes at low RMSD and the smooth transition to a new ΔG in the higher RMSD. These effects occur at different RMSD values for each mutant because they have different degrees of similarity to the available crystal structures. Please note that the single and double mutants had restraining walls acting at 5 and 4 Årespectively, while the triple mutant had no wall. The free energy profiles are thus extremely robust until we reach the RMSD region at which the walls start to act.

